# The diverging evolutionary history of opsin genes in Diptera

**DOI:** 10.1101/2020.06.29.177931

**Authors:** Roberto Feuda, Matthew Goulty, Nicola Zadra, Tiziana Gasparetti, Ezio Rosato, Nicola Segata, Annapaola Rizzoli, Davide Pisani, Lino Ometto, Omar Rota Stabelli

## Abstract

Opsin receptors mediate the visual process in animals and their evolutionary history can provide precious hints on the ecological factors that underpin their diversification. Here we mined the genomes of more than 60 Dipteran species and reconstructed the evolution of their opsin genes in a phylogenetic framework. Our phylogenies indicate that dipterans possess an ancestral set of five core opsins which have undergone several lineage-specific events including an independent expansion of low wavelength opsins in flies and mosquitoes and numerous family specific duplications and losses. Molecular evolutionary studies indicate that gene turnover rate, overall mutation rate, and site-specific selective pressure are higher in *Anopheles* than in *Drosophila*; we found signs of positive selection in both lineages, including events possibly associated with their peculiar behaviour. Our findings indicate an extremely variable pattern of opsin evolution in dipterans, showcasing how two similarly aged radiations - *Anopheles* and *Drosophila* - can be characterized by contrasting dynamics in the evolution of this gene family.

## Introduction

The ability to respond to specific visual stimuli is fundamental in defining patterns of behavioural activity, including predation and foraging in all animals. Opsins, expressed in photoreceptor cells, mediate light sensing in almost all living animals’ phyla (Feuda et al. 2012, 2016; Fleming et al. 2018, 2020). The modification of opsin repertoire (such as gene duplication or loss) and/or specific amino acid mutations in opsin genes(Feuda et al. 2016; Tierney et al. 2015, 2011; Sondhi et al. 2020) can confer the ability to adapt to new ecological niches.

Dipterans include approximately 10% of known animals, including iconic model organisms and several species of economic importance such as agricultural pests (e.g. fruit flies) and public health relevance such as vectors of infectious disease (e.g. mosquitoes) (White & Elson-Harris 1992; Attardo et al. 2014, 2019; Neafsey et al. 2015; Rota-Stabelli et al. 2020). Despite their importance and the clear link between opsins and sensory ecology (Feuda et al. 2016; Tierney et al. 2015, 2011; Sondhi et al. 2020), we lack a systematic understanding of the evolutionary history of these genes along the various lineages in Diptera. In this order, and in insects in general, the visual process is mediated by R-opsins classified accordingly to their maximum wavelength receptor capability: Long-Wavelength Sensitive (LWS) opsins can respond to green, Short Wavelength Sensitive (SWS) to blue, UV to ultraviolet light, and *Rh7* to a broad spectrum of light(Feuda et al. 2016; Sakai et al. 2017; Ni et al. 2017). Most of the knowledge we currently have on opsins in Diptera is based on *Drosophila melanogaster*, which is characterized by multiple paralogs of LWS (namely *Rh1* and *Rh*6), SWS (*Rh*2 and *Rh*5) and UV (*Rh3* and *Rh*4) (Sakai et al. 2017). Mosquitoes are characterised by a series of duplication events of LWS-*Rh*6 (Giraldo-Calderón et al. 2017), but whether these events are shared with other Diptera remains unclear. Furthermore, other opsin genes such arthropsins (that belongs to the R-opsins), the C and the RGR/GO have been identified in some insect groups(Almudi et al. 2020; Futahashi et al. 2015; Fleming et al. 2018), but their presence and eventual distribution in dipterans is ambiguous. As a consequence, it is unclear when dipteran opsins diversity evolved and it is difficult to study if and how they have played a role in their ecological flexibility.

To address these problems, we investigated the evolution of opsin genes in 61 Dipteran species sampled from ten different families, focusing on the model genera *Drosophila* and *Anopheles*. Our results substantially clarified the pattern of gene duplication of opsins genes in Diptera, identified several lineage-specific evolutionary events, and pointed toward events that might have contributed to define their ecology and behaviour.

## RESULTS AND DISCUSSION

### Dipterans have an ancient complement of five opsin genes and lineage specific expansions

Using a combination of BLAST, motif search and manual curation (see material and methods and Table S1), we identified a total of 528 opsins in 61 complete proteomes. We reconstructed their evolutionary affinities using Maximum Likelihood (Figure S1) and Bayesian (Figure S2) inference using the amino acid GTR-G model, which has been previously shown to fit opsins alignments better than other models (Feuda et al. 2012; Vocking et al. 2017). Both approaches (Figure 1, S1 and S2) revealed that there are at least eight opsin paralogous groups in Diptera, which we named using the *Drosophila* nomenclature as *Rh1-7* and C. We also found that arthropsins and RGR/GO opsins have been lost in Diptera. Furthermore, both methods recover a similar topology where LWS opsins (*Rh1, Rh2*, and *Rh6*) are monophyletic (PP=1 and BS= 100) and *Rh5* is the sister group to Rh3 plus Rh4 (PP=1 and BS=100). The position of *Rh7* is uncertain, with Maximum Likelihood favouring their position as sister group to all the remaining r-opsins (*BS=92*, Figure S1). Our tree of Figure 1 indicates several lineage-specific duplications and losses of opsin paralogs. To better understand their distribution in the various dipterans’ groups, we mapped their presence/absence into the Diptera phylogeny (Figure 2A-B). Results indicate that, starting from a common repertoire of five opsin genes (that we named *Rh1/2/6, Rh3/4, Rh5, Rh7* and *C* according to their phylogenetic relationships), this complement underwent a substantial change in a lineage-specific manner.

**Figure 1.**
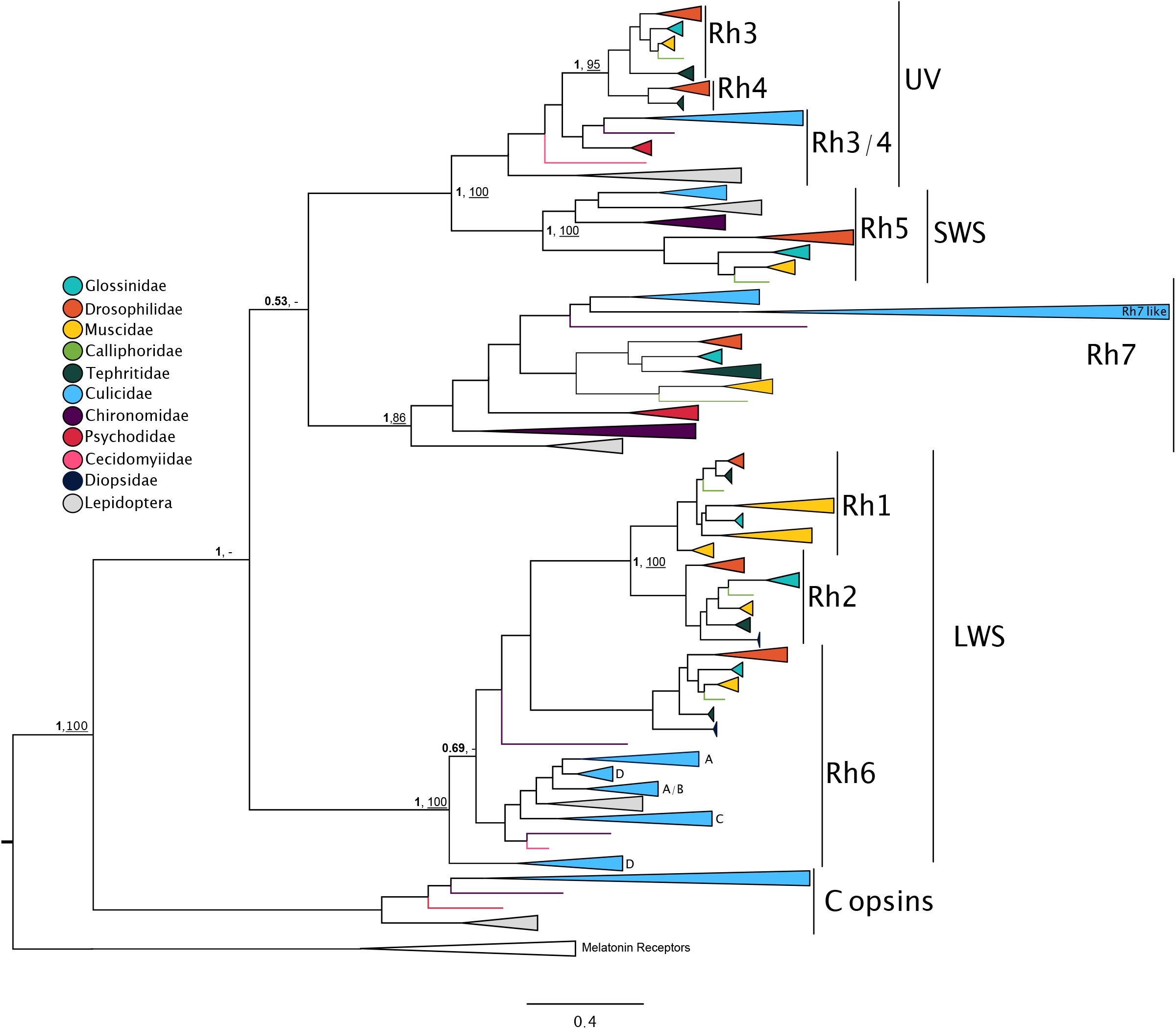
Phylogeny of 528 opsin from 61 dipteran species. Nodal support indicates the Bayesian Posterior Probability (in bold) and the UltraFast bootstrap (underlined). The colors indicate the different Diptera subgroups analyzed in the work.

**Figure 2.**
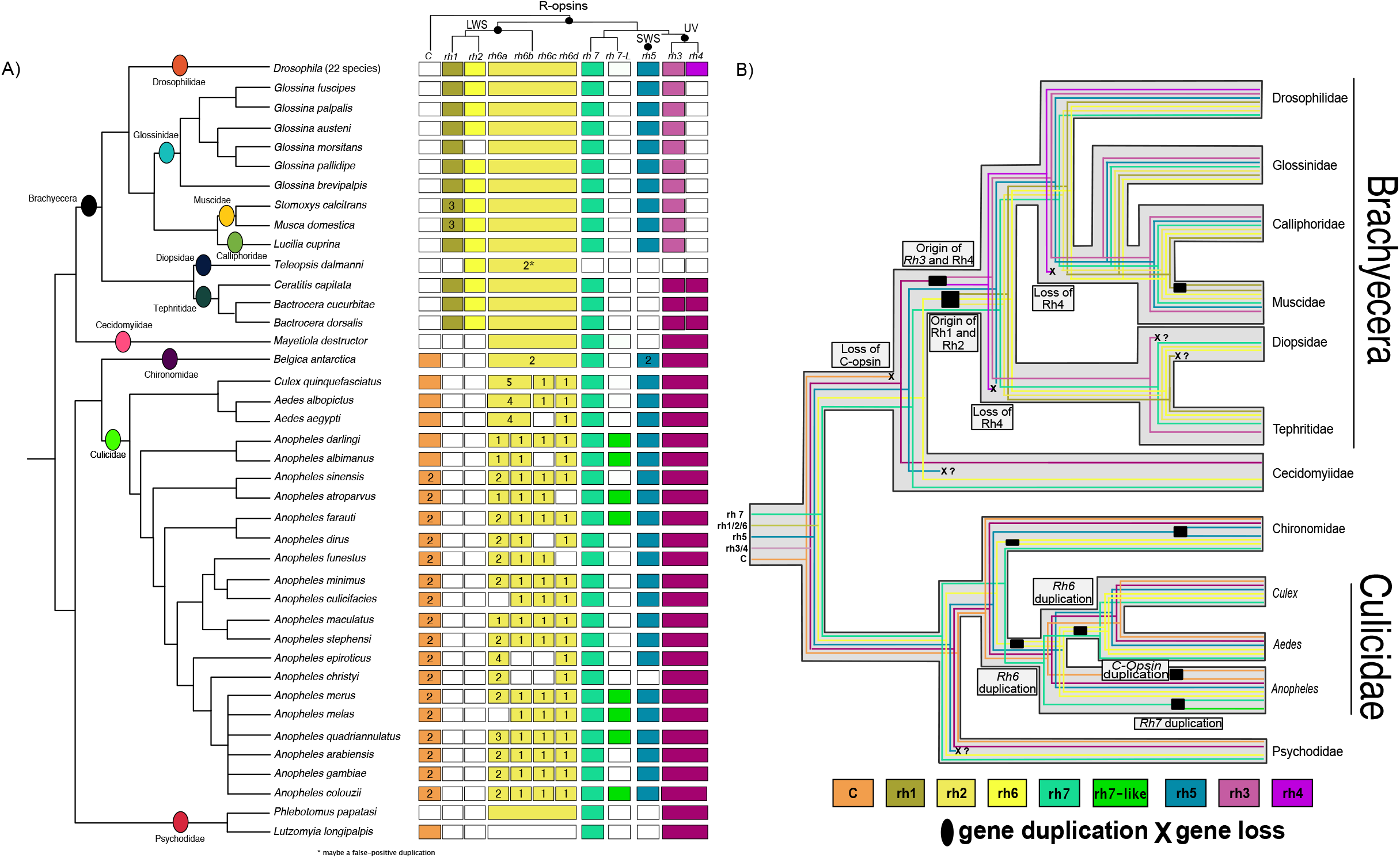
The phylogenomics of opsins in Diptera. **(A)** Opsin gene complement in the Diptera. Phylogenetic tree was obtained from to (Wiegmann et al. 2011). Gene nomenclature has been obtained from *Drosophila melanogaster*. Number indicates the numbers of genes identified; white boxes indicate that genes have not been found. **(B)** Synopsis of the patterns of opsin duplications and losses in Diptera subgroups. Lineage specific events are marked with a question mark if they were inferred from one single representative genome.

The opsins complement in the Brachycera (the clade comprising *Drosophila)* is highly derived compared to the ancestral condition. We confirm previous findings that c-opsin has been lost in all the Brachycera (Feuda et al. 2016)and can now demonstrate that four paralogs - *Rh1*, *Rh2, Rh3* and *Rh4* - are present only in this group. The shared presence of at least one duplication in *Drosophila*, tephritid fruit flies, muscidae house flies and *Glossina* tze-tze flies indicates that these duplications happened early in the Brachycera evolution from the ancestral Rh1/2/6 and Rh3/4 (Figure 2). We further observe various lineage specific events such as the loss of *Rh4* in the common ancestor of Glossinidae, Muscidae and Calliphoridae, duplications of *Rh1* in Muscidae, the loss of Rh2 in the tsetse fly *Glossina morsitans* (see also Attardo et al. 2019), and the loss of all opsins except for Rh2 and Rh6 in Diopside stalk-eyed flies. Interestingly, when we map the presence of introns (Table S2) in the different opsins, the results indicate that *Rh3* in all *Drosophila* species are intronless, suggesting their possible origin as retrotransposons.

In Culicidae mosquitoes (e.g. *Culex, Anopheles* and *Aedes)*, the repertoire of opsins is markedly different from the one observed in Brachycera (Figure 2). For example, eight out of the 19 *Anopheles* species have a divergent copy of the *Rh7* gene (Figure 1 and 2), whose phylogenetic distribution suggests that it was present in the ancestral *Anopheles* and secondarily lost in some species. We further found that *C-opsin* has been duplicated early in the *Anopheles* radiation. The most remarkable difference we observed between Brachycera and Culicidae is the impressive series of duplications of the *Rh6* gene in the latter, ranging from three copies in *Anopheles melas* and *Anopheles christyi* to seven in *Culex quinquefasciatus*. We identified four *Rh6* paralogs according to their relatedness (Figure 1) and distribution across the dipteran phylogeny (Figure 2), which we named *Rh6a*, *b*, *c* and *d*. These duplications have already been identified in three Culicidae species (Giraldo-Calderón et al. 2017), but our data indicate that this pattern is present in all the sampled Culicidae species, suggesting that at least two concomitant duplications happened in the Culicidae common ancestor, which were followed by additional lineage specific duplications. Despite a careful manual curation (see material and methods), we could not define the precise evolutionary relationships of these duplications within *Anopheles:* this is because some species lack well-assembled genomes and because of likely concerted evolution of paralogs in the same species. We found however that, similarly to *Rh3* in *Drosophila*, multiple *Rh6* paralogs lack introns (Table S2), suggesting that these newly evolved genes may have originated from a retrotransposition event.

The pattern of gene duplication summarized in Figure 2B allows for a substantial clarification of the opsin evolution in Diptera. Overall, our findings indicate that the opsin complement in Brachycera is highly derived compared to the ancestral condition in Culicidae, with the two groups having independently and differentially increased their repertoire of LWS opsins. Our detailed comparison *of Drosophila* and *Anopheles*, two genera for which we could analyse circa 20 genomes, showcased how the patterns of opsin evolution can dramatically differ in Diptera. While all *Drosophila* species have exactly the same opsin complement, indicating a frozen repertoire of opsin over circa 60 million years, the similarly aged *Anopheles* genus is characterised by an extremely plastic opsin repertoire as revealed by duplications and in some cases losses of *Rh6, Rh7* and C (Figure 2 and Table S1).

### The majority of newly evolved opsin genes contributed to the visual system

Our results so far indicate the Brachycera and Culicidae have divergent opsin complements with several lineage-specific events (Figure 2A and B). The question, therefore, arises whether these newly duplicated genes are expressed in photoreceptors cells and thus may be associated with a divergence and specialization of the visual system. In *D. melanogaster* there is ample evidence that all the opsin genes, including the newly duplicated intronless *Rh3*, are expressed and functional in photoreceptors cells and combinatorially define the different visual neural circuits(Courgeon & Desplan 2019). We further investigated the opsin expression in this species by mining single-cell RNA-seq data obtained from its entire optic lobe (Davis et al. 2020 and Figure S3). This data indicates that opsins expression is not restricted to the photoreceptors cells and that they contribute to different aspects of visual neural circuits. For example, the *Rh7* mRNA is detected in the lamina neurons L1-2 (Figure S3) that regulate motion. Furthermore, the function of the newly duplicated opsin genes in *D*. *melanogaster* may even not be restricted to the visual process: for instance, it has been recently proposed that *Rh1*, *Rh4* and *Rh7* are involved in chemo sensation(Leung & Montell 2017; Leung et al. 2020) suggesting a co-option of visual genes in different sensory pathways. In mosquitos, the information on opsin gene expression is instead scant. However, it is interesting to note that the R7 photoreceptors of *Aedes aegypti* may express, depending on their actual position in the retina, the LWS (*Rh6a-AAop2* or *Rh7-Aaop10*) the SWS *Aaop9* (*Rh5*) and the UV-(*Rh3*–*Aaop8*) opsins (Rocha et al. 2015). We further investigated the expression of opsin genes in *Anopheles gambiae* as reported in a microarray dataset (Baker et al. 2011): results indicate that *Rh6a*, *Rh3/4* and *Rh5* are statistically over-expressed in the head (Table S3), suggesting that they potentially play a role in the visual process. In conclusion, expression data indicates that several of the newly Brachyceran specific opsins genes as well as some of the mosquito specific *Rh6* copies are expressed in the photoreceptors and contribute to the function of the visual system.

### Opsins in mosquitoes and fruit flies underwent substantial divergent molecular evolution

Our results indicate that the opsin complement is highly divergent across the various Diptera families and that the majority of opsin genes are expressed in the photoreceptors and therefore involved with the visual system. This suggests that the different number of opsin paralogs may play a role in defining each species’ visual niche. However, adaptation to new niches might also be achieved by positive selections acting on single opsin genes. To test this, we investigated the pattern of molecular evolution in the 21 *Drosophila* and 19 *Anopheles* species included in the present study. To this aim, we produced manually curated opsin alignments for each paralog group and estimated the selective pressure that acted on these genes using the tool PAML (see material and methods for details). Importantly, the cross-group comparison is possible because both these two genus have a similar evolutionary history: both emerged in the Paleogene (between 100 and 30 mya according to Neafsey et al. 2015; Obbard et al. 2012) and have a similar number of generations per year (up to 10). We found that the differences between *Drosophila* and *Anopheles* are not restricted to the opsin repertoire alone but extend to the pattern of molecular evolution of the opsin genes. While these two groups show unusual signature of selection for a similar number of genes (shown in colour in Figure 4a; these are 23 and 26, respectively), in *Anopheles* we observe more events of site-specific positive selection (6 in *Anopheles* vs. 1 in *Drosophila*, magenta squares in Figure 4). A second difference concerns the rate of amino acid evolution (Figure S4), with an average *d*_N_ = 0.0118 in *Anopheles* compared to the *d*_N_ = 0.0073 in *Drosophila*, even though the two clades are characterized by a similar rate of synonymous nucleotide substitution (*d*_S_ = 0.2012 and *d*_S_ = 0.1969, respectively). This indicates that opsin genes are subject to different molecular constraints in the two groups, as supported by a slightly lower selective pressure in *Anopheles* than in *Drosophila* (Figure S5; overall *d*_N_/*d*_S_ = 0.0573 and *d*_N_/*d*_S_ = 0.0374, respectively). Combined with the positive selection tests, these results point to a much higher variability in selective pressure across opsin genes in *Anopheles*. These different evolutionary patterns are independent of data treatment: when regions with gaps are removed from the alignments (Figures S6, S7, S8) we observe lower substitution rates in *Anopheles* (because the orthologs in this genus are less constrained and accumulated more indels), but trends are consistent. Overall, our results indicate that in the genus *Anopheles* the opsin genes experienced a different evolutionary path and were subject to an accelerated rate of evolution compared to the *Drosophila* species; this is consistent and complementary with the more dynamic pattern of gene deletions/duplications we identified in *Anopheles*.

Current evidence indicates that in some cases it is possible to suggest a link connecting the variability in the evolution of the opsins across *Drosophila* species and ecological traits. For instance, the high heterogeneity in the selective pressure acting on *Rh7* (Figure S5), coupled with its fast evolution (i.e. high dN, Figure S4), may be associated to the diverse strategies adopted by the circadian clock in species with different ecology and latitudinal distribution (Menegazzi et al. 2017) and to the newly discovered taste abilities of this opsin in *D. melanogaster* (Leung et al. 2020; Leung & Montell 2017; see also supplementary materials for an extended discussion on the expression in *D. melanogaster*). We found that the agricultural pest *Drosophila suzukii* is characterized by an increased selective pressure affecting *Rh6* and *Rh7* (Figure 3). These changes are interesting from an applicative point of view, as they may be linked to the peculiar circadian activity (Hansen et al. 2019), colour recognition pattern (Little et al. 2019), and even gustatory preferences(Crava et al. 2016; Leung et al. 2020) deriving from the evolution of this species’ new ecological lifestyle. In *Anopheles* the function of opsins is less understood when compared to *Drosophila* (Montell & Zwiebel 2016). However, mosquito species may be active at different periods of the day or night depending on the genus, when light is characterized by different wavelength composition while other sensory stimuli are not predicted to vary substantially in a circadian fashion (e.g. CO_2_, smell). The unique capacity of the opsins to tune their maximum absorbance to specific light conditions might thus have had a role in these ecological differences (Jenkins & Muskavitch 2015). Indeed, we hypothesise that the high variability in the selective pressure affecting *Rh7* and C-opsin in *Anopheles* species (Figure S5) may be linked to differences in their adaptation to light. It is our hope that these and other patterns of molecular evolution (Figure 3) as well as the many genetic novelties (Figure 2) we have identified will inspire neurophysiological studies aimed at improving our understanding of flies and mosquitoes visual biology and improve their management practices.

**Figure 3.**
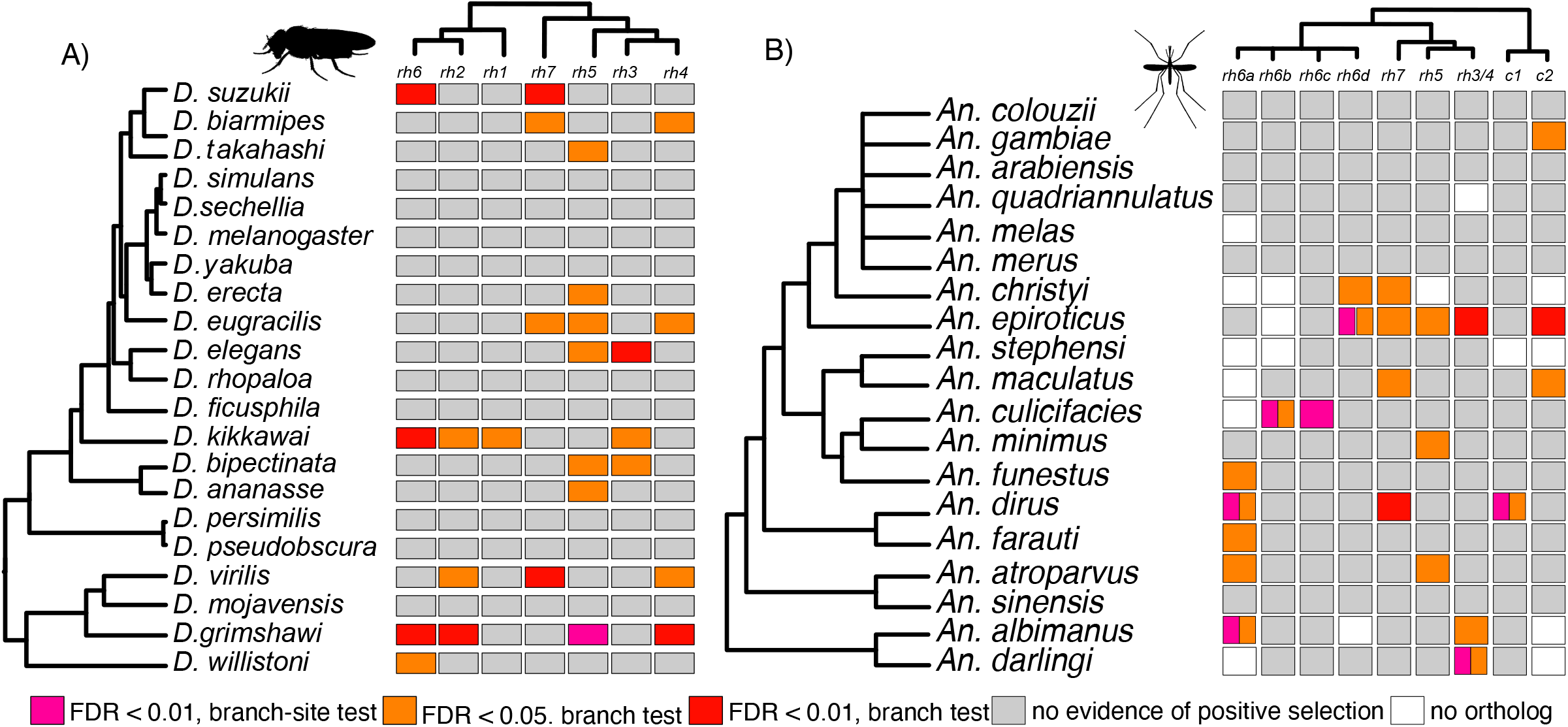
Pattern of positive selection for the opsins genes in *Drosophila* (A) and *Anopheles* (B). (A) Comparison of ω for each paralogue across all *Drosophila* (B) and *Anopheles* species. These analyses were performed including the columns containing gaps.

## CONCLUSIONS

We have characterized in detail the evolutionary history of the opsin genes in ten dipteran families, with a focus on the fine scaled molecular evolution of model organisms *Drosophila* and *Anopheles*. Overall, we found that, starting from a core of five opsins genes, different groups of dipterans underwent distinct patterns of deletions-duplications (Figure 1 and 2). For example, our results reveal that the opsins complement in *Drosophila* (and in general in Brachycera) has diverged considerably from its ancestral state, with several genes present only in this group (e.g. *Rh1* and *2*). This in turn suggests that *Drosophila* is not a very suitable model to infer the ancestral opsins function (see also Pisani et al. 2020). Our data also indicate that in mosquitos the opsin complement is much more dynamic, particularly concerning *Rh6* and *Rh7*, which suggests it might have had an important role in the ecological success of mosquitoes, such as the invasion of urban dwellings by *Anopheles*. Although our analyses have revealed that the *Anopheles* and the *Drosophila* lineages have experienced a similar amount of positive selection affecting the evolution of their opsins (Figure 3), we have found more instances of site-specific selective pressure and faster evolutionary rates in *Anopheles* (figure S4 and figure S5). More broadly, the evolution of the opsins in *Drosophila* and *Anopheles* showcase how diverging patterns of genes evolution can characterize two chronologically similar radiation episodes. Overall, our findings reveal an extremely variable pattern of opsin evolution in dipterans, a result that may be exploited to improve our understanding of their biology and control.

## MATERIALS AND METHODS

### Opsins identification

We downloaded 61 predicted proteomes from 11 Diptera families (see table S1). To identify the opsins genes we employed a combination of BLAST and motif research similarly to Feuda et al. (2016). In brief the sequences from (Feuda et al. 2012, 2016; Ramirez et al. 2016) were used to mine every genome. From this analysis every gene with an e-value below 10^-10^ was retained as putative opsins genes and subject to motive research using Prosite (Sigrist et al. 2013) and annotation using BLASTP against the Uniprot90 Database. To be considered an opsin one of the two conditions was sufficient: a sequence must contain a retinal binding domain or have an opsin as first hit in the BLAST search. Using this approach and after a preliminary manual annotation we identified 528 opsins genes see table S1. Alignment and trees are available on bitbucket (https://bitbucket.org/Feuda-lab/opsin_diptera/src/master/).

### Manual curation

The dataset obtained was eventually manually curated. First, we looked for putative false duplication in the tree. Where we detected two sequences from the same species which were clustered together with high similarity in our phylogenies (a putative recent duplication), we checked if they share the same locus, and when this condition was verified we deleted the shorter of the sequences. These false duplications have likely occurred because two opsins proteins were predicted from the same gene with two different start codons or intron boundaries. Secondly, we checked for missing data. We selected sequences that missed part of the opsin protein, and, where possible, we retrieved the missing data using BLAST (tblastn) on the assembled genomes. Moreover, we looked for unexpected opsins losses to assess whether the missing genes were true losses or artifacts. In some cases, we found the missing gene in the genome of interest and the sequence was added manually to the alignment. For some mosquito species we lacked well-assembled genomes and therefore accurate gene models. This may have caused misrepresentation in the exact number of *Rh6* copies in each lineage and blurred the fine-scale pattern of duplications/losses. To mitigate the impacts of this, we specifically validated manually the *Rh6* genes in *Anopheles* species. Using such an approach allowed us to increase the length of many orthologs but most importantly, we could detect instances of false positives. For example, we identified cases where putative duplicated contigs or allele variants from heterozygotes genomes could be mistaken for species-specific duplications.

### Gene structure characterization

To infer the molecular mechanisms underlying the origin of new opsin genes we investigated the presence of introns in the 61 Diptera species under study using Vector base (Giraldo-Calderón et al. 2015), Ensembl (Yates et al. 2020), and in some cases by manual curation. Our results indicate that several genes, including *Rh3* in all *Drosophila* species and multiple *Rh6* paralogs in some mosquitos, lack introns (Table S2).

### Gene annotation

The full gene region of each of the 7 opsins in Drosophila was inspected in the flybase genome browser for the presence of introns, and each intron presence and length was mapped in Table S3. To assess significant events of intron length variation, we assumed a normal distribution for the length of each of the introns in the sample and highlighted as significant, those intorns that were outside, or at the edge of two times the standard deviation for that intron, excluding from the average the target intron.

### Phylogenetic analysis

To identify the phylogenetic relationships between the opsins genes we performed a phylogenetic analysis using Bayesian and maximum likelihood inferences. The opsins data set was then merged to a subsample of the insect dataset of Feuda et al. (2016), sequences were aligned using MAFFT (Katoh et al. 2002) and default parameters. The maximum likelihood tree was performed using IQTree 2.0 (Nguyen et al. 2015) under GTR-G4 model. The Bayesian tree was performed using Phylobayes-MPI (Lartillot et al. 2013) under GTR-G4 model and nodal support was estimated using Posterior Probability (PP).

### Positive selection

In this case the coding sequence for case each opsin subgroup (*Rh1, Rh2, Rh3, Rh4, Rh5, Rh6, Rh7*, and *C*) were aligned separately for the 21 *Drosophila* and the 19 *Anopheles* species using PRANK (Loytynoja & Goldman 2008)codon model, which produces less false-positives in positive selection analysis(Markova-Raina & Petrov 2011). Each alignment was manually curated to avoid spurious divergence signals that may have biased the subsequent analyses and we generated two sets of alignments, one using all sites and a second where all the regions containing gaps were removed. We inferred the level of selective pressure acting on each of the 7 Drosophila opsins (dataset 3) using PAML 4.7 (Yang 2007). Rates of synonymous (*d*_S_) and nonsynonymous substitution (*d*_N_), as well as their ratio ω = *d*_N_/*d*_S_ (which measures levels of selective pressure acting on a gene), were estimated by the “free-ratio” model using the tree topology from (Ometto et al. 2013). To allow cross opsin-gene comparisons in *Anopheles*, in this analysis, alignments included only those species present in all opsin alignments. Heterogeneity in the selective pressure was inferred using a branch-test to compare the likelihood of a single ω model across branches (model = 0 and NS sites = 0) versus one assuming two distinct ω, one for each terminal branch, one at a time (i.e. for each *Drosophila* and *Anopheles* species in their respective datasets), and another for rest of the tree. To further identify the occurrence of positive selection on specific sites we employed the branch-site test (branch-site model A, test 2; model = 2 and N_S_ sites = 2; null model has parameters fix_ ω = 1, ω = 1; the positive selection model fix_ ω = 0, ω = 1, with each species set as foreground species in separate analysis, see above). Both tests were estimated using either the whole alignment (clean=0) or removing parts of the alignment where one or more sequences contained a gap (clean=1). For both tests, we tested twice the difference between the log-likelihood of the two models using a χ2 test with 1 degree of freedom. To account for multiple testing, we estimated the false discovery rate (FDR) of each test using the q-value approach (Storey 2002) implemented in R (R Development Core Team, 2009).

---

**Figure S1**. Maximum likelihood tree of 528 opsins genes under GTR-G. Nodal support indicates Ultra-Fast Bootstrap value.

**Figure S2**. Bayesian tree of 528 opsins genes under GTR-G. Nodal support indicates Bayesian posterior probability value.

**Figure S3**. **Analysis of *Rh* genes in *Drosophila* sensory system the data were obtained from** (Davis et al. 2020) focusing of photoreceptors cells.

**Figure S4. Rate of protein evolution (non-synonymous substitution, *d*_N_) across opsins phylogenies in *Drosophila* (A) and *Anopheles* (B)**. Different letters identify significant statistical differences at adjusted *P* < 0.05 according to a Tukey’s HSD (honestly significant difference) multiple comparison test. Median and quantiles are shown as grey lines for each gene.

**Figure S5. Pattern of positive selection (d_N_/d_S_) for the opsins genes in *Drosophila* (A) and *Anopheles* (B).** These analyses were performed including all alignment position. Different letters identify significant statistical differences at adjusted *P* < 0.05 according to a Tukey’s HSD (honestly significant difference) multiple comparison test. Median and quantiles are shown as grey lines for each gene.

**Figure S6. Summary of pattern of positive selection for the opsins genes in *Drosophila* and *Anopheles*. (A)** Comparison of ω for each paralogue across all *Drosophila* **(B)** and *Anopheles* species. These analyses were performed excluding columns containing gaps.

**Figure S7. Rate of protein evolution (non-synonymous substitution, dN) across opsins phylogenies in *Drosophila* (A) and *Anopheles* (B)**. These analyses were performed excluding columns containing gaps. Different letters identify significant statistical differences at adjusted *P* < 0.05 according to a Tukey’s HSD (honestly significant difference) multiple comparison test. Median and quantiles are shown as grey lines for each gene.

**Figure S8. Pattern of positive selection for the opsins genes in *Drosophila* (A) and *Anopheles* (B).** These analyses were performed excluding the columns containing gaps. Different letters identify significant statistical differences at adjusted *P* < 0.05 according to a Tukey’s HSD (honestly significant difference) multiple comparison test. Median and quantiles are shown as grey lines for each gene.

**Table S1.** Opsin complement in 61 Diptera species

**Table S2. Intron distribution in 61 Diptera species.** Red squares indicate opsins that are intronless. When multiple copies of the genes are present the number in parentheses indicates the intronless opsins.

**Table S3.** Opsins expression obtained from Baker et al. 2011.

## ACKNOWLEDGMENTS

This study was supported by a Royal Society University Research Fellowship (UF160226) to R.F.

## Supplementary materials

### Extended discussion

#### On the expression of Drosophila opsins in light sensory systems

There is ample evidence that all the opsin genes we have studied are expressed and functional in *Drosophila melanogaster* (reviewed in Courgeon & Desplan 2019). A recent study examining single-cell RNA-seq in the optic lobe provides further, and at times unexpected, findings about the expression of opsins in this species (Davis et al. 2020). The study confirms that the majority of opsin genes, including the intronless *Rh3*, are expressed in retina photoreceptors (Figure S3). Conversely, *Rh7* is not only detected in photoreceptors cells; instead, it is also expressed in neurons in the lamina, medulla and lobula plate (Figure S3). This result puts to rest a controversy concerning the expression of *Rh7*, which was detected in the retina by only one of two antibodies being developed(Kistenpfennig 2012; Ni et al. 2017). It is interesting to note that Rh7 is expressed in the lamina neurons L1, L2 and L3 that receive direct synaptic connections from R1-R6 photoreceptors (Figure S3). L1 and L2 are part of two distinct pathways regulating motion detection, whereas L3 is involved in a pathway regulating color vision (Silies et al. 2014). The expression of opsins in neurons (which qualifies them as photoreceptor cells) that also receive input from other photoreceptor cells (which qualifies them as interneurons) is a common logic in light detecting/processing circuits. This circuit organization suggests that the canonical photoreceptors are also implicated in circadian entrainment by light. Many experiments have shown such an assumption to be true. However, in *D. melanogaster* an additional twist adds to the story. The R8 photoreceptor cells expressing Rh6, which correspond to *ca*. 65% of all R8 cells in the retina (see below), are particularly important for light entrainment, likely via excitatory cholinergic neurotransmission (Alejevski et al. 2019). The latter is specific to this group of photoreceptor neurons and is present in all retinal photoreceptor cells in *D. melanogaster*, in addition to the standard inhibitory histaminergenic transmission (Davis et al. 2020). However, a dual photoreceptor interneuron function is a recurrent theme, not only in circadian light pathways for the entrainment of the circadian clock, but also in visual processing, as exemplified by the color opponency mechanism. Light carries two types of information, “luminance” that is used for the perception of form and motion and “colour” that is used to add contrast and detail to forms. Once light has been absorbed by a photopigment, like an opsin, changes in the electrical properties of photoreceptor cells tell the nervous system that a luminous stimulus of certain intensity has been perceived. To interpret colour, additionally the nervous system compares the response of different photoreceptor cells with different spectral sensitivities. To improve discrimination, such a comparison is antagonistic and must rely on interactions between at least two different channels (i.e. photoreceptor types). In *Drosophila* it starts with the reciprocal inhibition of the R7 and R8 photoreceptors that are part of the same ommatidium(Nguyen et al. 2015).

A surprising finding of Davies et al (2020) is the unexpected ‘expansion’ (compared to what known to date) of opsin types in photoreceptors. In many sensory systems the “one neuron – one receptor” rule allows the unambiguous detection of information across the whole sensory field. For instance, in *Drosophila* a combination of stochastic processes, gene expression exclusion and coordinated development results in the production of “yellow” and “pale” ommatidia approximately in 2:1 ratio and randomly distributed across the retina. The expression of opsins is strictly coordinated: yR7-R8 express *Rh4-Rh6* whereas pR7-R8 express *Rh3-Rh5*. However, about 10% of “yellow” ommatidia break the “one neuron – one receptor” rule as their R7 express both *Rh3* and *Rh4*, while their R8 express *Rh6*. Consistently, these ommatidia are located in the dorsal half of the eye (Mazzoni et al. 2008). If the findings by Davies et al. (2020) were confirmed, the situation might be even more complicated. The R7 photoreceptors (y+p) seem to express *Rh2, Rh3, Rh4, Rh5, Rh6* and the yR8 allegedly express *Rh4*, *Rh5*, *Rh6* (Figure S3). Although the co-expression of opsins was not examined, it would seem a distinct possibility if these observations were validated at the protein level.

Co-expression of opsins is the rule rather than the exception in mosquitoes. In *Aedes aegypti*, the retina can be divided into four domains, dorsal, central, ventral stripe and ventral, according to the opsins expresses in the R7 and R8 photoreceptors. The R7 photoreceptors express the LWS *Aaop2* + SWS *Aaop9* in the dorsal and ventral stripe regions whereas they express the LWS *Aaop10* + the UV *Aaop8* in the central and ventral domains. This broad spectral sensitivity is thought to maximize photon capture, albeit at the expense of color discrimination, a perfect compromise considering these organisms are active under dim light conditions (Rocha et al. 2015)

#### More on the involvement of Drosophila opsins to non-light sensations

Recently it has emerged that opsins have sensory functions unrelated to light sensing. Although extra-retinal opsins have been documented in vertebrates for quite some time, all non-visual functions previously identified depended on photoreception nevertheless (Leung & Montell 2017). In the last few years work on *Drosophila melanogaster* has documented that opsins contribute to thermosensation in larvae (*Rh1, Rh5, Rh6)*, to proprioception during larva locomotion (*Rh1, Rh6)*, to adult hearing (Rh5, Rh6 and possibly Rh3, Rh4) and to adult bitter taste discrimination (*Rh1, Rh4, Rh7*)(Sokabe et al. 2016; Zanini et al. 2018; Senthilan et al. 2012; Leung et al. 2020). All these functions are independent from photoreception, which begs the question on how these sensory modalities avoid interference from light. This has been clarified for proprioception and hearing (that depend on mechanoreceptors) and taste (that depends on gustatory receptor neurons), the neurons responsible for those sensory modalities are able to traffic opsins that are not bound to their chromophore; in all other cells known to date – including those where opsins function as a thermosensor – the chromophore serves as a molecular chaperone required for the export of the protein from the endoplasmic reticulum (Leung et al. 2020). These physiological differences and the evolutionary history of the different opsins involved suggest that these light-independent functions in *Drosophila* are derived (Pisani et al. 2020).

