## Supplementary figures and images for "The diverging evolutionary history of opsin genes in Diptera"

### FigureS1

FiguresS1

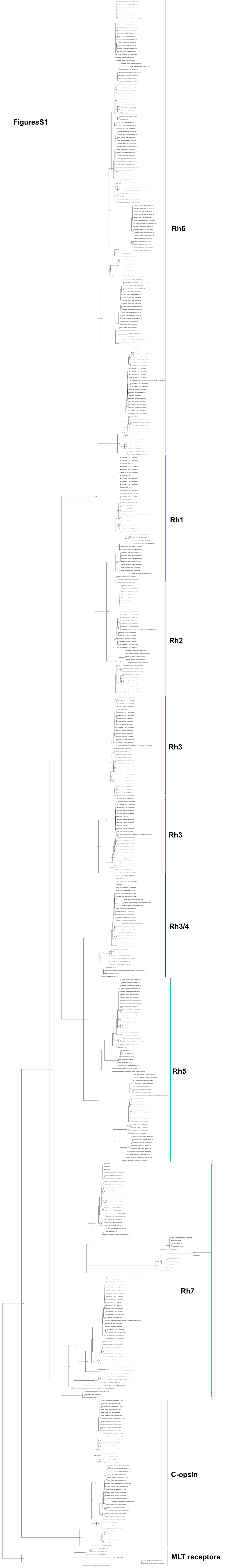

### FigureS2

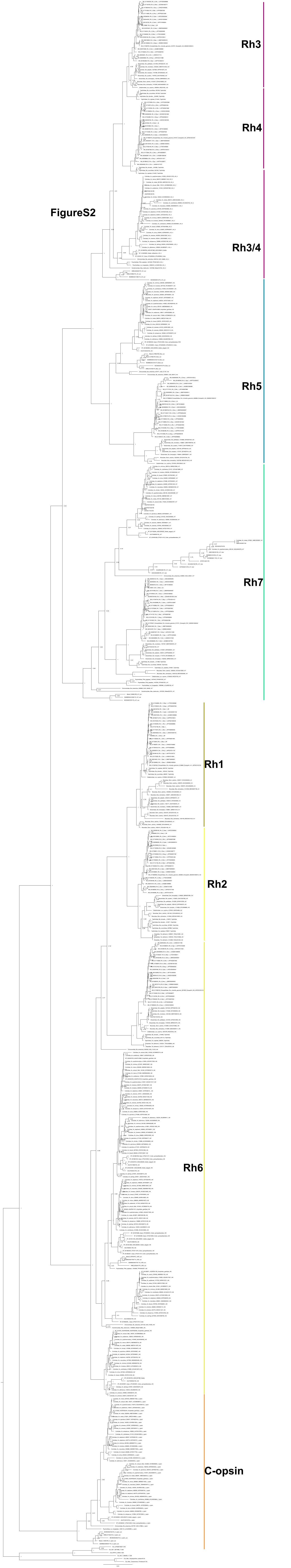


FigureS2

### FigureS3

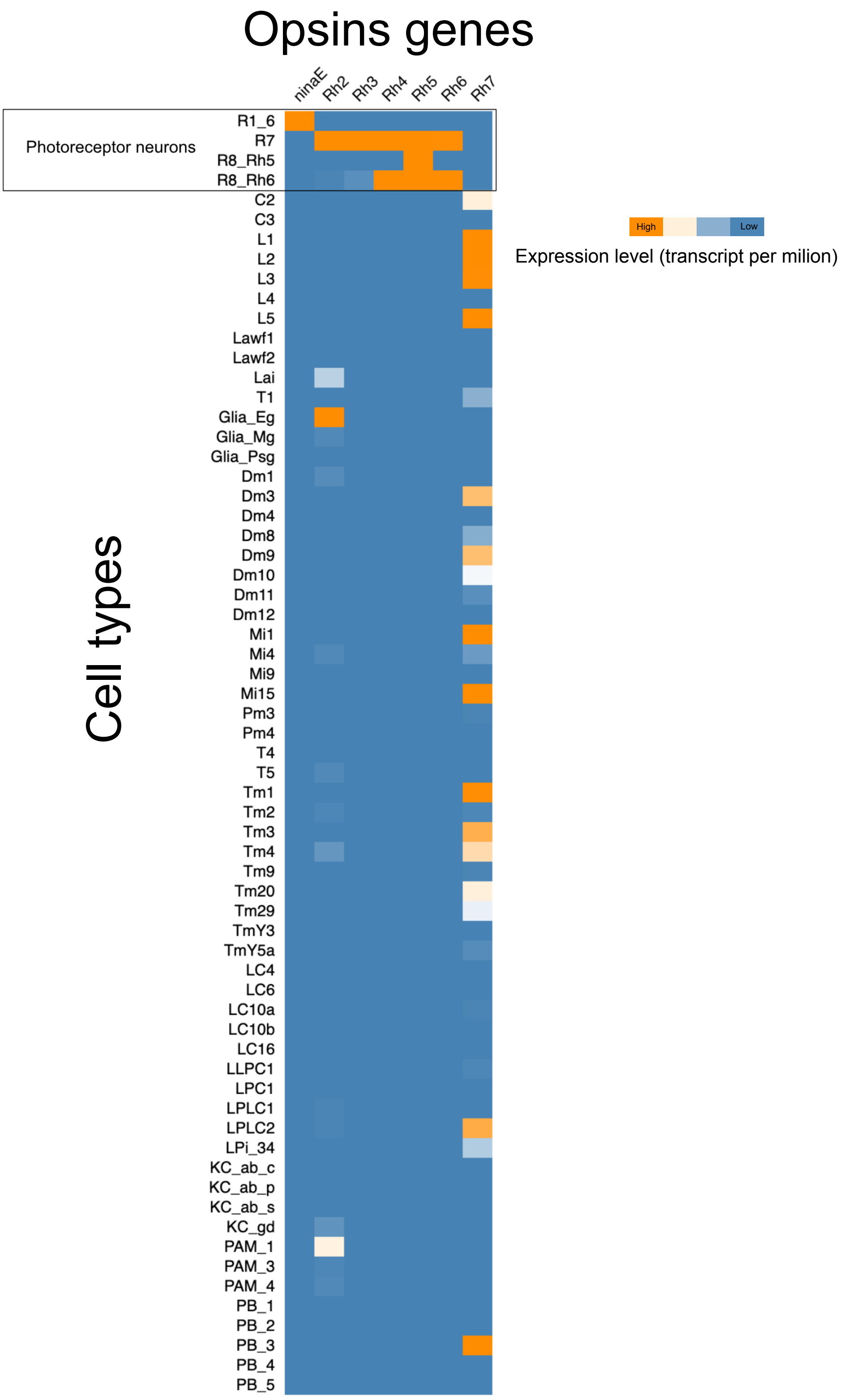

### FigureS4

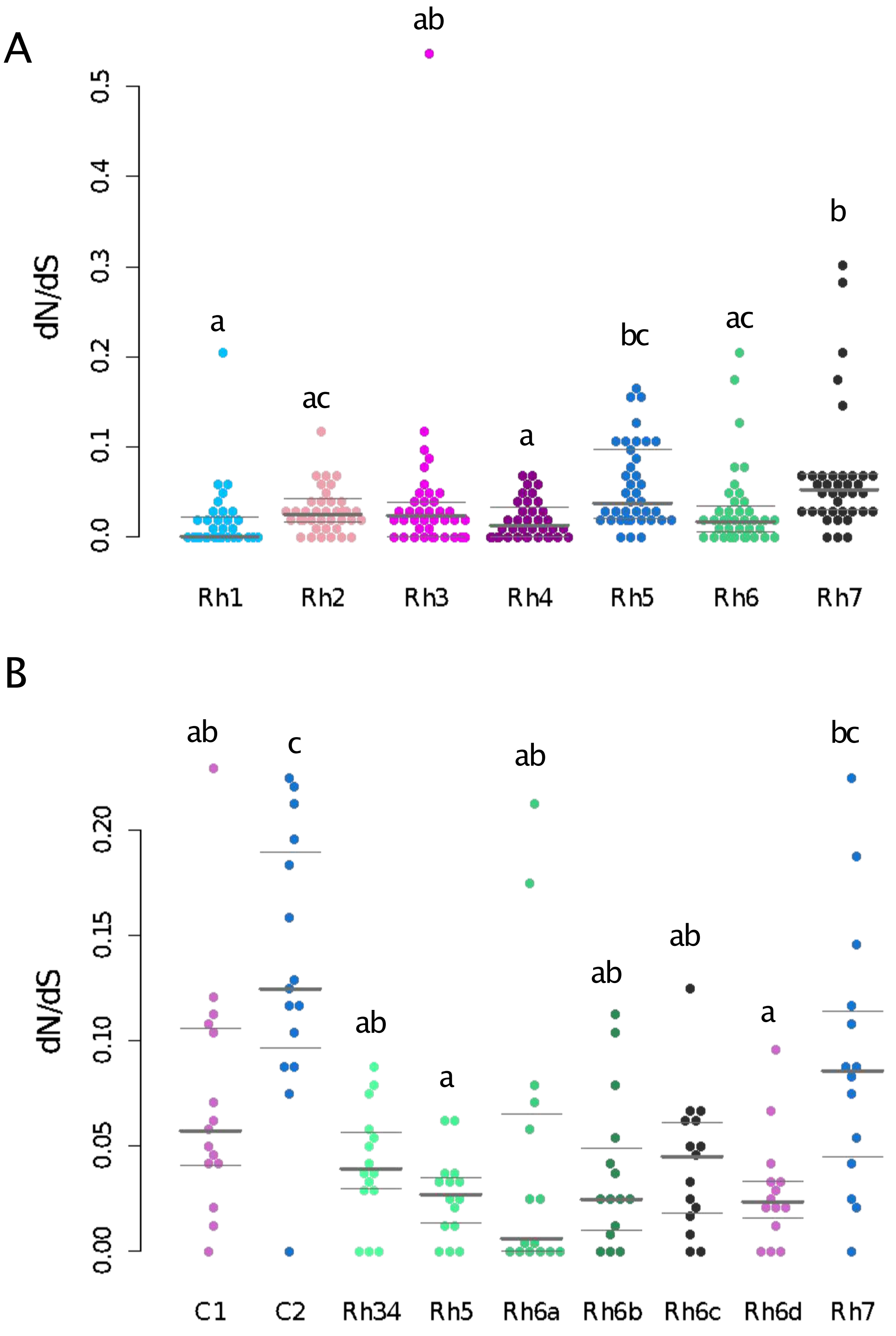

### FigureS5

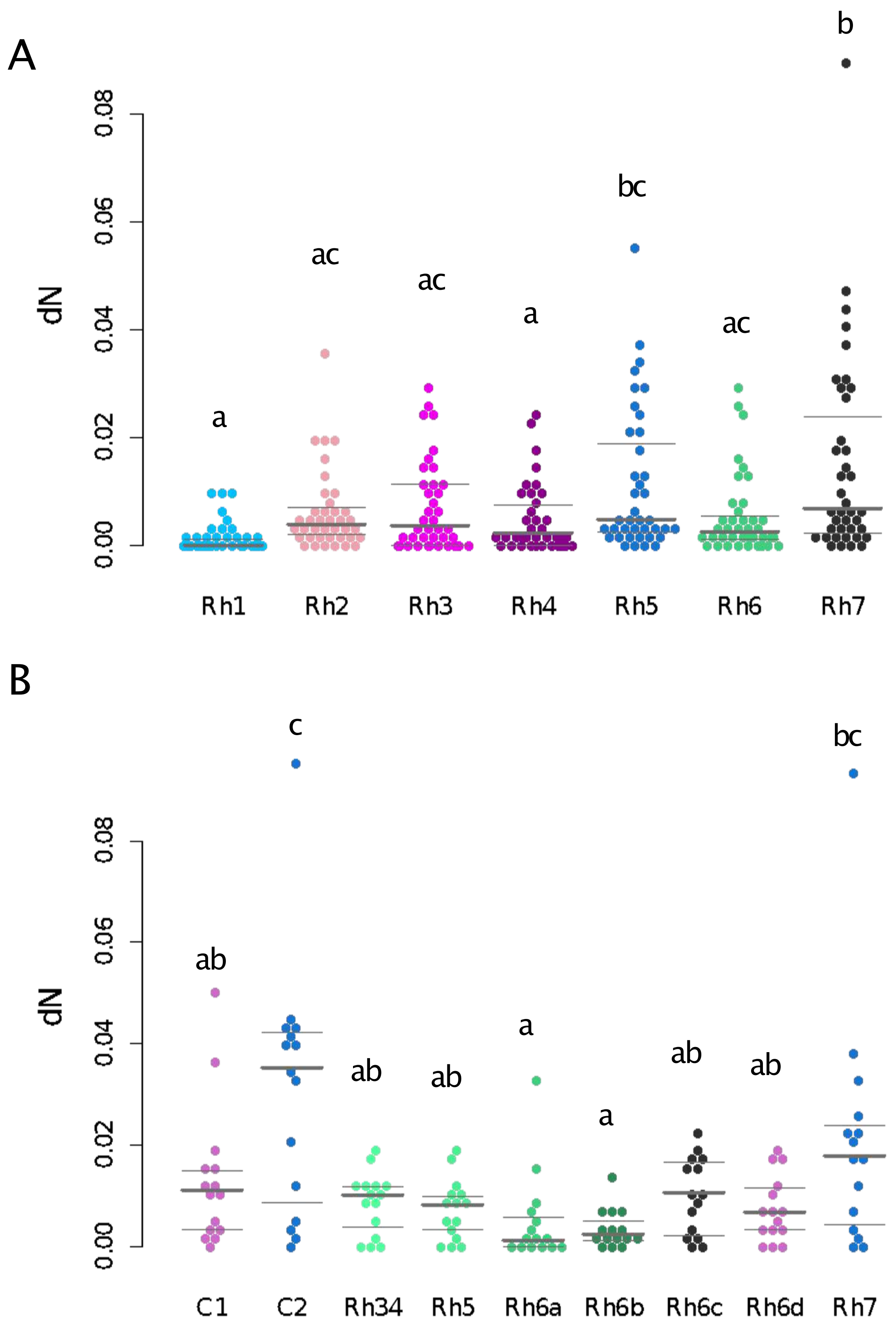

### FigureS6

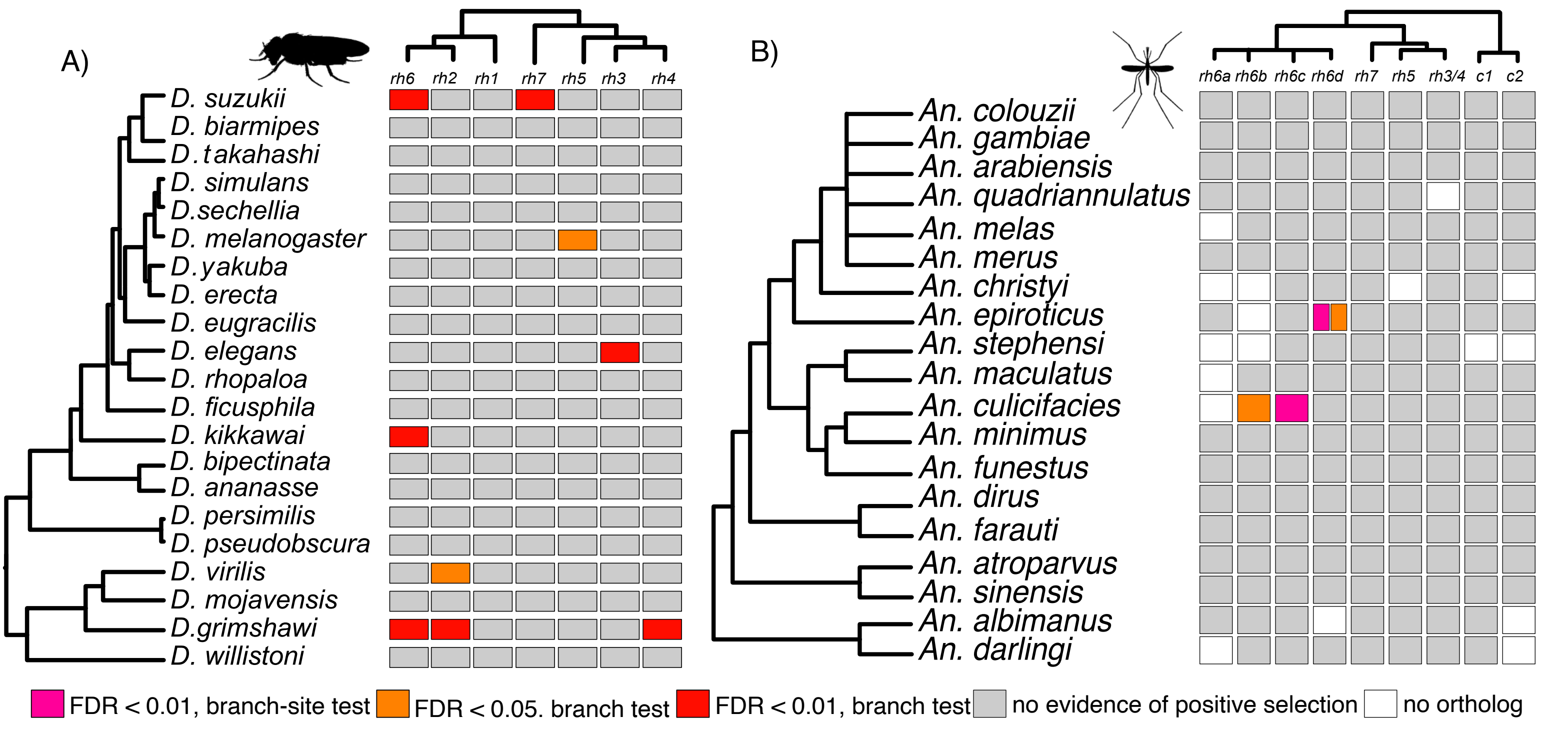

### FigureS7

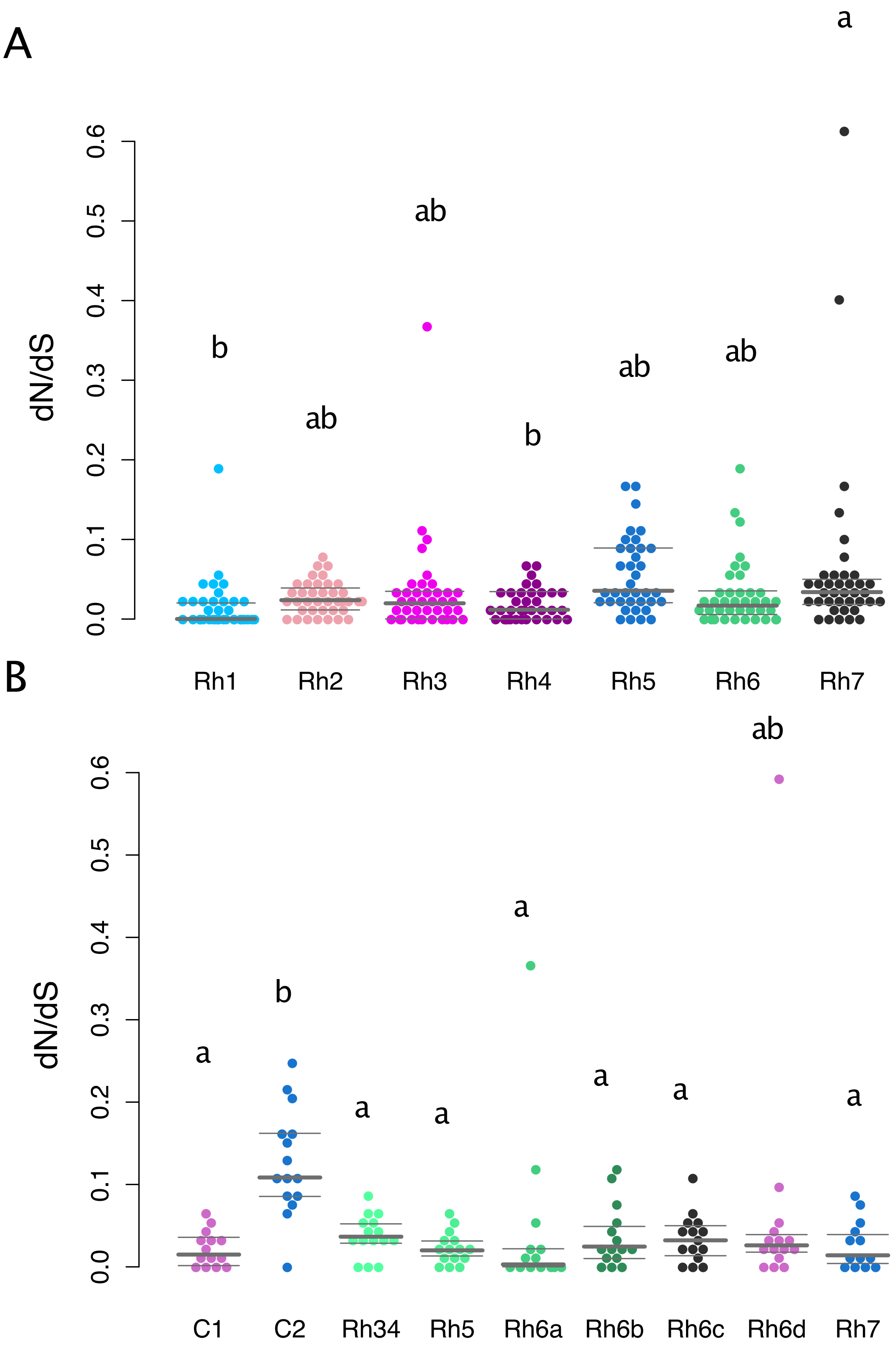

### FigureS8

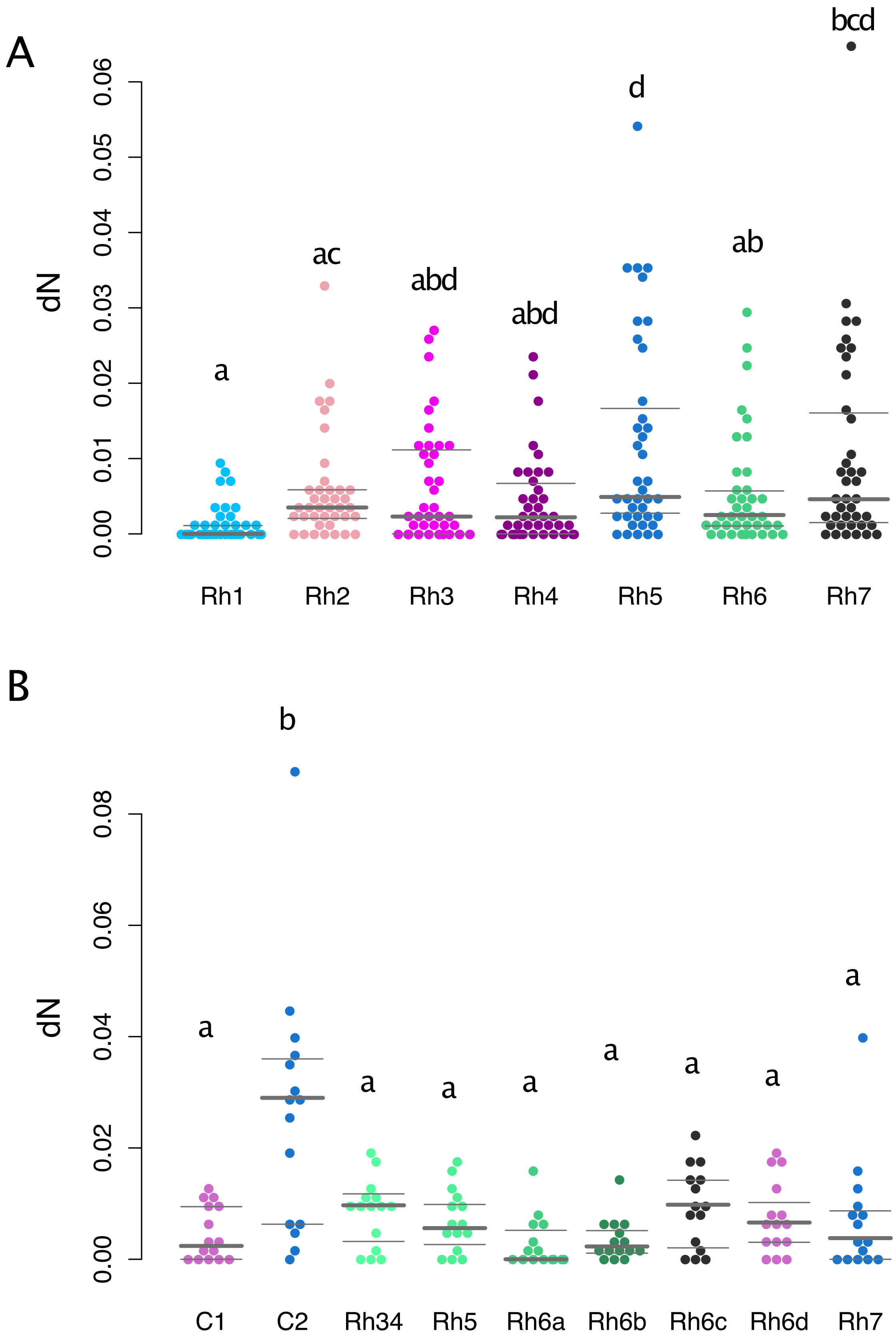
